# Concurrent predictive and prospective strategies in a simple visuomotor task

**DOI:** 10.1101/2024.05.15.594355

**Authors:** Inmaculada Márquez, Mario Treviño

## Abstract

Interception, a fundamental visuomotor skill for activities such as driving and sports, involves two main strategies: predictive, anticipating the target’s trajectory, and prospective, actively tracking and adjusting movement. Experimentally controlled factors could potentially influence the relative usage of these strategies. We designed a visuomotor task to probe the relationship between target predictability and interception strategies. We manipulated stimulus predictability through controlled adjustments of external forces, altering the target’s trajectory. We also manipulated the availability of perceptual information by introducing spatial occlusion at specific parts of the visual field. Our observations indicate that decreased target variability promoted predictive interception, whereas increased variability prompted a shift toward prospective strategies. Notably, hand-catching trajectories exhibited increased curvature in response to changes in target variability, whereas eye trajectories displayed a relatively consistent curvature across trials. Similarly, heightened target variability resulted in delayed onset of hand movements while showing no discernible alterations in the onset of eye movements. Thus, gaze position was a poor predictor of hand position, highlighting distinct adaptive patterns for hand and eye movements in response to task unpredictability. Finally, participants exhibited consistent interception strategies within and across sessions, highlighting their differences and preferences for predictive or prospective strategies. These results reveal a dynamic interplay between target predictability and interception, suggesting a flexible combination of both approaches. Examining how humans integrate sensory information, plan, and execute movements provides a unique opportunity to characterize predictive and prospective interception strategies in dynamic, real-world scenarios.

## 1 INTRODUCTION

Our ability to predict and adapt to the environment is a fundamental skill for survival. Such predictive capacity, rooted in our brain’s ability to anticipate sensory information, shapes and controls our actions. Furthermore, according to a new theory called ‘predictive processing’, reality, as we experience it, is mainly built from our predictions (Keller & Mrsic-Flogel, 2018; Clark, 2023). Employing tasks that involve intercepting moving objects offers an ideal setting for examining predictive control strategies. The neural substrates for interception encompass various brain regions (*e.g*., primary visual cortex, posterior parietal cortex, dorsal anterior cingulate cortex, primary, premotor, and supplementary motor cortex, basal ganglia, and cerebellum) responsible for processing target motion, timing motor responses, and updating internal models (Zago *et al*., 2009). Research also suggests multiple routes for visuospatial information to reach frontal regions (*e.g*., dorsolateral prefrontal cortex, responsible for working memory, decision-making, and planning complex actions; anterior cingulate cortex, monitors performance and adjusts behavior based on feedback), potentially influencing interception mechanisms (Roussy *et al*., 2021).

Individuals executing predictive interception strategies plan their movements by anticipating future events. In other words, this approach predicts the future location of the target by considering its position, trajectory, and speed. Predictive control is particularly effective when the target maintains a constant speed and sufficient time is available for interception (Beek et al., 2003; Eggert et al., 2005). On the other hand, prospective control prioritizes real-time movement adjustments as additional information about the target’s path becomes available (Dessing *et al*., 2009; Merchant *et al*., 2009). It relies on continuous guidance from perceptual information and is particularly useful when the target exhibits unpredictable motion or when the time window for interception is constrained. Thus, prospective control requires precise sensorimotor coordination and quick responses to changing target motion. It is important to emphasize that, although predictive and prospective sources of information can coexist within a particular context, they serve distinct purposes in guiding behavior. In other words, each strategy exerts unique effects on movement patterns (Muller & Abernethy, 2006).

Predictive control often results in more planned and less variable movements (*i.e*., less curvature), while prospective control generates gradual movement modulation, increasing adjustments as actions unfold (*i.e*., more curvature; (Panchuk & Vickers, 2009; Zago et al., 2009; Katsumata & Russell, 2012; Ledouit et al., 2013; Zhao & Warren, 2015). Some authors have suggested that using a predictive or prospective strategy depends on external elements, such as target motion, but also on internal elements, like cognitive differences across individuals (Lee *et al*., 1997; Fooken *et al*., 2016; Treviño *et al*., 2023). Therefore, various factors, including the quality of sensory information, task complexity, and individual skill and experience, could influence the relative usage of these two strategies (Eggert *et al*., 2005; Dessing *et al*., 2009; Merchant *et al*., 2009).

While some researchers have suggested the necessity of incorporating predictive and prospective control to explain a variety of interceptive actions (Panchuk & Vickers, 2009), the question arises as to whether these strategies can operate concurrently within a single task. How do humans use predictive and prospective strategies to intercept moving targets under varying target predictability and visibility levels? We developed a simple task where participants used a mouse cursor to catch a falling dot on a screen. Our experiments aimed to characterize how task conditions influenced the utilization of predictive and prospective interception strategies. Our central hypothesis was that enhancing the predictability of the target’s motion would encourage the adoption of predictive strategies, whereas introducing uncertainty, such as incorporating random forces in the target’s horizontal direction, would favor prospective strategies. To further examine this idea, we manipulated the availability of perceptual information by introducing spatial occlusion in specific parts of the visual field. This manipulation prompted participants to dynamically update their strategies whenever they gained visual access to the falling target (Panchuk & Vickers, 2009). Our results demonstrate the coexistence of predictive and prospective strategies within a single interception task. The utilization of these strategies was intricately tied to the predictability and visibility of the falling target. Our results also evidenced a weak correlation between gaze position and manual (control of the optical mouse/cursor) response, pointing to distinct roles for the ocular and motor systems in the interception process. Finally, there were enduring individual variations in the utilization of predictive and prospective strategies across different conditions and sessions, emphasizing the personalized nature of strategy selection within this visuomotor task. The coexistence of predictive and prospective strategies highlights the flexibility in movement control, indicating that humans can continuously employ a versatile combination of both approaches. Switching between these strategies should be crucial for successful performance in real-world situations.

## 2 MATERIALS AND METHODS

### 2.1 Participants

Our study involved 61 adult participants, comprising 32 women and 29 men, aged between 19 and 22 years, with a mode age of 21. None of the participants had been previously diagnosed with psychiatric, neurological, or neurodevelopmental disorders, and they had no history of substance use. Most participants were right-handed (≥ 95%, indicating their dominant hand for writing) and had either normal vision or vision corrected to normal, as confirmed by a Snellen test. For the study procedures, participants received written instructions for questionnaire completion and task execution. We also collected demographic information with these questionnaires. Participation was entirely voluntary and not associated with monetary incentives. We used quota sampling to enlist undergraduate students from our university. Participants were assigned to the experimental groups to ensure a gender-balanced composition. This approach maintained equal and independent probabilities for participating in the assigned tasks. The sample size was pre-established using the G*Power program (Faul *et al*., 2007), ensuring adequate group numbers to replicate experimental findings (Trevino, 2014). We obtained informed consent from all participants, and all procedures adhered to non-invasive methods while fully complying with our country’s local guidelines and regulations. Our research received ethical approval from the Instituto de Neurociencias ethics committee, Universidad de Guadalajara, México, under the reference ET102021-330.

### 2.2 Visuomotor task

The visual task involved a falling white dot with a fixed size (0.3° of visual angle) on a 27-inch computer monitor with a resolution of 1920 × 1080 pixels and a refresh rate of 60 Hz. The dot was initially placed at the upper center of the screen and remained still for 300 milliseconds before initiating its motion. Upon release, the dot acquired an initial velocity along the y-axis and moved horizontally in either the left or right direction of the monitor. In the variability-free mode, the dot’s motion was governed by three primary factors: its initial horizontal velocity (negative for leftward motion, positive for rightward motion), a constant gravitational component inducing downward acceleration along the y-axis, and air friction introducing acceleration opposing the direction of motion. A small blue rectangular object (width of 2° visual angle and height of 0.3° visual angle) was located at the bottom of the screen, under the user’s control through an optical mouse. The aim of the task was for the user to capture the descending dot by adjusting the horizontal position of the blue rectangular object (vertical position of the blue rectangle always fixed to the bottom of the screen). This horizontal movement could be executed preemptively, before the dot’s descent, or in real-time as the dot descended. In other words, participants could choose their preferred approach, introducing a strategic element to the task (**Figure 1**). We provided auditory feedback in the task through earphones, using short tones with pure frequencies (10 kHz for successful trials and 1 kHz for unsuccessful trials (Treviño *et al*., 2023).

**FIGURE 1.**
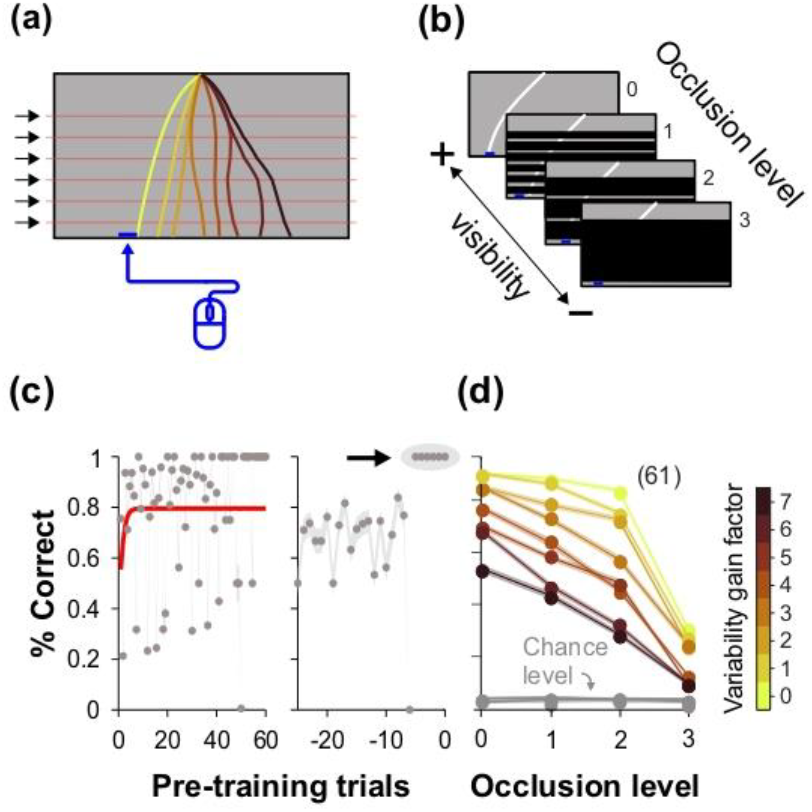
Falling dot task to study predictive and prospective human behavior. (a) Cartoon of the experimental setup. Participants controlled a blue rectangle’s horizontal movement at the screen’s bottom using an optical mouse while seated. The objective was to catch a falling dot, allowing users to choose between preemptive or real-time adjustments, introducing a strategic element to the task. (b) The falling dot’s trajectory was influenced by six adjustable forces at equidistant imaginary lines and intermediate heights (variability. These forces perturbed the horizontal path, modified the speed, and affected the landing position, allowing controlled adjustments (gain factor from 0 to 1). (c) The task allowed for the introduction of a mask segment obscuring the visibility of the falling target below a specified height (visibility). The target’s initial trajectory and speed were observable for a vertical distance equivalent to 20 times its size. Two modifications were made to the ‘whole mask,’ maintaining its dimensions. The first variant included two horizontal slits, each with a height of 4 times the target size, positioned to intermittently reveal the falling target. The second variant featured five slits and six intermediate masks symmetrically positioned within the mask, creating a spectrum of visual occlusion from no visibility to full visibility. (d) Different levels of variability and visibility were tested independently, and their combinations were systematically permuted throughout the experiments. This design allowed us to map participants’ responses along the variability and visibility axes. (e) Participants were pretrained until achieving five consecutive correct catching trials to ensure task proficiency, which was accomplished in less than 100 trials. Participants had an average learning asymptote of approximately 80% (Trevino, 2020). (f) Catching performance against masking conditions (x-axis; colorbar on the right-hand side represents target stimulus variability). Performance declined with increasing variability and masking conditions. Number of participants in parenthesis.

To manipulate the predictability of the falling target, we’ve made the next procedures. In the initial step, we simulated the trajectory of the falling point and adjusted the initial speed parameters, falling acceleration, and friction acceleration. These adjustments ensured that the particle’s descent consistently positioned it at the midpoint of the monitor screen’s (left or right half), with a trial duration of 2.3 seconds (**Supplementary Figure 1a**). Therefore, without additional perturbing forces (*i.e*., in the variability-free mode), the target would naturally descend to one of these fixed positions. We introduced six additional forces at equidistant imaginary lines and intermediate heights (**Figure 1b**). These forces were applied to the descending particle’s horizontal trajectory, perturbing its path, modifying its horizontal speed, and influencing its ultimate landing position. The direction (left or right) and intensity of these forces were intentionally adjustable to exert varying degrees of impact on the particle’s trajectory. We performed computer simulations to determine the maximum range of force magnitudes and directions, ultimately selecting a combination of random forces that resulted in final target positions within the screen boundaries (**Supplementary Figure 1b**). Subsequently, we introduced a virtual gain factor, adjustable from 0 to 1, enabling systematic and precise modifications to control the impact of these external forces (**Supplementary Figure 1c**).

To manipulate the available sensory information, we’ve masked segments of the target’s trajectory, occluding its visibility. The mask had a defined height, allowing participants to observe the target’s initial trajectory (and speed) for a vertical distance equivalent to 20 times the size of the target (**Figure 1c**). Below this specified height, the mask was activated, making the particle invisible to the participant until it reached the bottom of the screen. We introduced two modifications to the ‘full mask’ while maintaining its dimensions (full mask = occlusion level 3). The first variant featured two horizontal slits, each with a height of 4 times the size of the target, equivalent to 1.2° of visual angle. These slits were strategically positioned within the mask to offer intermittent visibility of the falling target as it could be visualized through them. The slits were symmetrically positioned within the mask to ensure accurate alignment (mask 2 = occlusion level 2). The second variant featured five slits (*i.e*., six intermediate masks), each of the same size as the previous. These slits were symmetrically positioned within the mask (mask 1 = occlusion level 1; **Figure 1c**). The idea behind using a full mask and its variants with increasing numbers of slits was to create a system covering a spectrum for visual occlusion (*i.e*., occlusion levels 0-3), ranging from complete visibility to no visibility, gradually decreasing the information about the falling target. In other words, with this design, a higher number of slits offered a more extensive and detailed view of the target’s falling trajectory, providing richer visual cues for interception. Conversely, a lower number of slits reduced the available visual information, introducing a degree of uncertainty into the interception process. At the far end of this spectrum, the full mask served as a visual barrier, blocking all information about the falling target beyond the initial visible portion. This experimental design allowed for a controlled exploration of the impact of varying levels of visual occlusion on participants’ ability to intercept the target. We randomly shuffled the masking conditions during the experimental sessions, mitigating potential order effects and minimizing the influence of any systematic biases that might arise from using a fixed sequence of conditions (Herrera & Trevino, 2015). In a task variation, we introduced color coding for the falling dot, representing negative horizontal speeds with red and positive speeds with green. The color intensity corresponded directly to the speed magnitude. This manipulation gave participants a visual cue about each trial, offering useful *a priori* information on the falling dot’s motion and aiding anticipation of its final landing side. We limited the experimental sessions to an hour to ensure participants remained engaged and focused.

### 2.3 Eye tracking

We employed an eye-tracking device to investigate the oculomotor control of successful interception. A high-precision commercial eye tracker (Tobii, Stockholm, Sweden) recorded binocular movements at a sampling rate of 60 Hz. Participants sat at a comfortable distance of 60 cm, while we fixed the device to the lower part of the monitor. We aligned the eye tracker headbox with the participants’ eye level and minimized head movements using a chin holder with a forehead rest. The two cameras of the tracker captured stereo images of both eyes, providing data on eye gaze, position, and pupil diameter. We maintained consistent room luminance during the experiments and programmed calibration routines to ensure accurate eye tracking during our experimental sessions (**Supplementary Figure 2**). The initial routine was a standard 4-point calibration conducted directly through the eye tracker software. We custom-designed a second routine in MATLAB with the assistance of the Psychophysics Toolbox, placing four fixation dots at the periphery. We segmented the experimental sessions into three blocks, incorporating 5-minute breaks between each block, and conducted calibration before each experimental block to ensure accurate tracking throughout the study. The eye tracker also recorded binocular pupil diameters under an average illuminance of approximately 100 lux (measured at a 60 cm distance from the monitor). To elicit the pupillary light reflex, participants underwent a pre-trial calibration routine involving six cycles of alternating contrast conditions (black and white full screens) repeated four times. We addressed missing data and blink artifacts through linear interpolation. All oculomotor and pupillary responses were processed using custom MATLAB routines.

### 2.4 Analysis

We continuously recorded the dot’s motion parameters using custom software developed in MATLAB R2022a, integrating the Psychophysics Toolbox extensions (PTB-3). We quantified the number of successful catches and computed the corresponding averaged probabilities. To contextualize these empirical values, we also determined the probability of a catch occurring purely by chance (Treviño *et al*., 2023). This was accomplished through a randomized process for each trial, using a uniform distribution of the user’s position along the screen’s horizontal axis. We conducted our analysis by focusing on the timing and spatial precision of both hand and ocular movements. As mentioned, the manual response data corresponded to the participants’ recorded actions as they manipulated the horizontal position of a small blue rectangle on the screen using an optical mouse. In some experiments, we recorded participants’ gaze positions to monitor their eye movements throughout each falling trajectory. Such dual measurements allowed us to characterize manual and visual elements associated with our task.

To investigate the interception strategy employed in each trial, we analyzed the optical mouse position (manual response) and eye-tracking data to characterize the participants’ focus on the target’s current and future positions. We calculated the squared Euclidean distances between the participants’ current position and gaze to the current and future target positions. This analysis used a defined time window spanning 25 forward frames, equivalent to approximately 416 milliseconds. We have justified the rationale for employing these metrics previously (Treviño *et al*., 2023). In evaluating the effectiveness of predictive and prospective strategies, we compared positive and negative squared distance differences relative to the one observed with the actual positions (*i.e*., those observed in present time). These differences should be negative for predictive strategies, implying the participant’s ability to anticipate the target’s trajectory. In contrast, these differences should be positive for future target locations than the current position in the case of prospective strategies. We measured the cumulative curvature of both manual and gaze movements by determining the magnitude of the curvature vector traced through all the points along each trajectory. We also examined the timing of participants’ movements, establishing thresholds of 30°/s and 50°/s to define the onset of manual and saccadic eye movements, respectively. We used empirical analysis to validate the manual threshold by calculating traces’ root mean square (RMS) before and after movement initiation. The threshold for eye tracking data, established through previous studies to differentiate rapid from slower eye movements, ensured that only rapid and ‘ballistic’ ocular movements were considered in the onset analysis (Klein & Ettinger, 2019).

### 2.5 Statistical analyses

We conducted a multivariate analysis of variance (MANOVA) to evaluate the effects of target predictability and visibility on the manual and gaze responses. Additionally, we conducted a Repeated-Measures (RM) variation of the Kruskal-Wallis test tests to assess the similarity of medians across visual occlusion levels. We also computed the Euclidean distance between the manual and gaze positions and the current and future target positions to characterize the interception strategies. We calculated the curvature and onset of the manual and gaze trajectories to distinguish between predictive and prospective control. We used Pearson correlation coefficients to assess the reliability of the test-retest and the correspondence between gaze position and manual response. We report group data using mean values and their corresponding standard error of the mean (S.E.M.). Statistical significance was set at *P* ≤ 0.05. The number of participants included in the analysis for each task is specified within the corresponding figure panels, enclosed in parentheses.

## 3 RESULTS

### 3.1 Visuomotor Task for Studying Predictive and Prospective Interception Behavior

We modified a visuomotor task (Treviño *et al*., 2023) to explore the interplay between predictive and prospective interception behaviors. Here, participants were required to catch a falling white dot projected on a computer monitor while retaining all the other main elements of the original task. Participants used an optical mouse to move a blue rectangle positioned at the bottom of the screen to intercept the falling dot. Initial speed, gravitational acceleration, and air friction (*i.e*., negative acceleration) influenced the motion of the falling dot. We introduced additional forces, activated at imaginary equidistant lines, to perturb the dot’s horizontal trajectory, impacting its final landing position on the bottom of the screen (**Figure 1a**). The task also involved occluding visual access to the falling dot, incorporating masks with multiple slits for intermittent visibility (**Figure 1b**). We have created four mask types to investigate the influence of visual occlusion on participants’ interception strategies. From lowest to highest: type 0, the “No Mask” condition consisted of trials without masks, offering an unobstructed view of the falling target. In additional mask types, all featuring the same height, participants could observe the initial trajectory of the target for a distance equivalent to 20 times its size. Type 1 incorporated a mask with five equidistant slits, providing intermittent visibility of the target’s trajectory. Type 2 utilized a mask with only two horizontal slits (with the same height as those for type 2 mask), resulting in less visibility of the target’s trajectory compared to type 1. Finally, type 3, the “Full Mask” condition, completely obstructed the visual field, rendering the target invisible until it reached the bottom of the screen (**Figure 1b**). In sum, the objective of the task was to capture a falling target under various visual conditions, introducing a strategic element when solving each trial. We controlled trial unpredictability through gain factor adjustments applied to the external forces acting on the falling target, while concurrently investigating the impact of visual information on participants’ interception abilities by manipulating masking conditions.

With this task, our objective was to explore how participants adjusted their interception strategies in relation to our manipulation of unpredictability and visual access. We hypothesized that individuals relying exclusively on prospective strategies might encounter difficulties as the target’s visibility decreased. Conversely, those transitioning toward predictive strategies could potentially maintain or even improve performance in conditions of reduced visibility. We started our experiments by introducing participants to a pre-training phase to ensure task comprehension. Participants were required to achieve five consecutive correct catches as part of the manual response analysis. Additional training trials were provided until this criterion was met (**Figure 1c**). Subsequently, by implementing multiple permutations, we assessed participants’ performance in the task across eight variability levels (0-7) and four occlusion levels (0-3) (**Figure 1d**). As expected, there was a reduction in manual interceptive performance as the occlusion level increased (average performance: mask type 0: 80.31% ± 0.18%; mask type 1: 69.70% ± 2.24%; mask type 2: 57.53% ± 2.49%; mask type 3: 19.83% ± 1.62%; *n* = 61), and as the target trajectory variability increased due to the influence of external forces (Several-sample repeated measures tests, *F* = 1.48×10^4^, *P* < 0.05 for all groups; w. effects across all parameters, MANOVA, *F*_6,950_= 55.13, *P* < 0.0001, confirmed with Multiple linear regression, *F*_3,8380_ = 7331.1, *P* < 0.05; **Figure 1d**). Hence, by considering the influence of external forces and visual occlusion on participant performance, this task provides a powerful platform for investigating interception strategies. At the beginning of the trials, participants should use a prospective strategy, tracking the initial phase of the falling particle trajectory. However, as the trajectory changes with the target’s movement and the introduction of external forces, participants may adapt, potentially transitioning from prospective to predictive strategies as the variability and visibility of the target’s trajectory change.

### 3.2 Interplay of target predictability, visibility, and interception strategies in our visuomotor task

In interception tasks, the nature of the target’s motion influences task difficulty, with increased predictability facilitating successful performance. Two main interception strategies employ predictive and prospective control (Mrotek & Soechting, 2007). Predictive interception is employed when dealing with predictable targets, where the interceptor relies on knowledge of the target’s current state and their own movements to anticipate the target’s future position. This strategy involves predicting the target’s upcoming location based on its past and current positions, typically applied in conditions characterized by simple and predictable target motion or when information is sporadic. On the other hand, prospective interception is employed for more unpredictably moving targets, demanding continuous updates. In this case, the interceptor relies on real-time information about the target’s speed and direction to deduce its arrival at the next immediate location. In principle, the choice between these strategies is linked to the level of target predictability, with reduced predictability favoring prospective approaches and heightened predictability promoting predictive strategies. Moreover, partial masking may shift a prospective strategy towards a more predictive one by limiting continuous perceptual information, prompting reliance on prior knowledge and anticipation of the target’s future position. Therefore, both predictive and prospective strategies involve making predictions about a target’s future position, but they differ in terms of the type of information they use and the situations in which they are most effective. Importantly, these strategies are not mutually exclusive, and their distinction is not always clear-cut.

In our main task design, we manipulated both target predictability and visibility. We examined participants’ manual catching responses (the optical mouse movements) to characterize interception strategies, considering the target’s present and future positions. Tracking the distance to the target’s future positions over time is valuable for distinguishing between predictive and prospective interception behaviors (de la Malla *et al*., 2019). Therefore, we developed an algorithm to compute the Euclidean distance between the mouse position (manual response) and current and future target locations, utilizing data from our experiments. We calculated forward traces within a time window of 25 forward frames, approximately 416 ms. In this context, a predictive manual strategy would be indicated by a decrease in distance relative to future target locations. Conversely, for a prospective manual strategy, the distance should consistently be greater for future target locations than for the current one. Each subpanel from **Figure 2a** displays the Euclidean distances over the last 100 frames (approximately 1600 ms). before the falling point reached the bottom of the screen. In other words, these traces are aligned to the moment when the falling dot contacted the bottom of the screen, depicted at the right extreme of each panel. To facilitate comprehension of our analysis, we illustrated the traces as the difference between future distances (depicted with darker shades of green) and the current one (blue horizontal trace). In this representation, when green traces exhibit a downward trend (negative differences), it indicates predictive behavior since the manual position is closer to the anticipated final location of the target. Conversely, an upward trend (positive differences) indicates prospective strategies, suggesting manual adjustments following changes in target position. **Figure 2a** illustrates participants’ adaptive responses to variability in target trajectory (columns) and visual access (rows). Reduced variability in the target trajectory resulted in catching trajectories dominated by predictive control. Conversely, with higher target variability, strategies became more prospective (Repeated measures ANOVA test, area under the curve, AUC, below forward traces; *F*_7,327_ = 44.3, *P* < 0.005; *n* = 41). These results demonstrate the coexistence of predictive and prospective control in our task.

**FIGURE 2.**
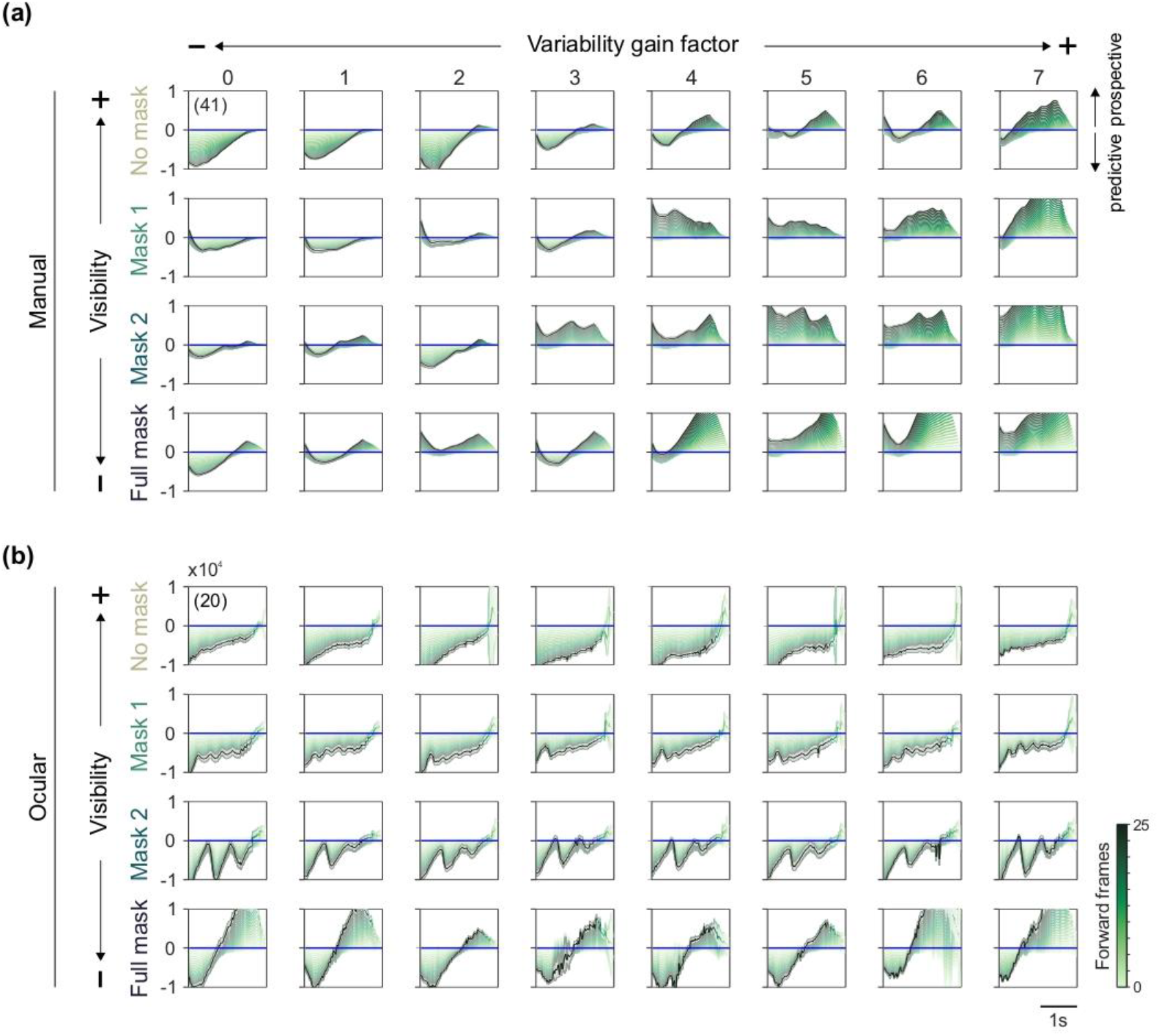
Coexisting predictive and prospective strategies in the falling dot task. Manual (a) and eye-tracking (c) data were analyzed to explore participants’ behavior in relation to the falling target’s current and future positions. We calculated Euclidean distances between user’s positions and current and future target positions using a 25-frame time window (∼416 ms). To facilitate visualization, we illustrate the differences between future distances (darker shades of green) and the current one (blue line). A downward trend (negative difference) involves predictive behavior, where the position aligns closer to the anticipated final target location. An upward trend (positive difference) suggests prospective strategies, indicating adjustments in response to changes in the target position. (b, d) Colormaps of the same traces shown in panels (a) and (c). This representation helps to visualize the predictive (blue) or prospective (red) nature of the interceptive strategy. Number of participants indicated in parentheses.

In conditions demanding rapid adjustments, individuals may lean towards prospective control based on continuous visual information (Panchuk & Vickers, 2009). In less challenging conditions, however, individuals might employ a mixture of predictive and prospective interception strategies (Fooken *et al*., 2016). For all these cases, ocular movements play a critical role in interceptive control by enabling the tracking of the target and estimation of distance, speed, and direction of travel. This information is essential for planning and executing successful interception movements. Therefore, we employed an eye-tracking system to explore the relationship between gaze patterns and the falling target. Following calibration and standardization on obtaining the eye-tracking data (**Supplementary Figure 2**), we conducted experiments depicted in **Figure 2b**, presenting the outcomes of the Euclidean distance-to-target analysis based on gaze data. These traces reveal that ocular positions employed predictive strategies (Repeated measures ANOVA test, *F*_7,159_ = 3.7, *P* < 0.005; *n* = 20). Notably, although the responses in the presence of type 2 masks confirm a gaze pattern dominated by predictive processing, the gaze quickly transitioned towards a prospective pattern when the target became visible through the mask’s slits. In other words, these rapid fluctuations in eye-tracking distance coincide with instances where the user observed the target through the slits and engaged in a prospective strategy, minimizing the distance to anticipated target positions.

### 3.3 Distinguishing predictive from prospective control using curvature and timing

Our methods differentiating between predictive and prospective control indicate that both strategies coexist in our visuomotor task. Predictive behavior manifested with falling trajectories exhibiting minimal variability, while increased variability promoted adopting prospective control. An additional approach to distinguish predictive from prospective control involves examining the curvature of the catching trajectories (Vickers, 1996; Fooken *et al*., 2021). In prospective control, an interception trajectory typically exhibits high curvature, reflecting constant adjustments to intercept the target. On the other hand, predictive control often results in a more linear interception trajectory, indicating straightforward movement toward the target’s predicted location. We assessed the sensitivity of manual and eye-tracking responses to stimulus unpredictability by examining the curvature of the manual (only the horizontal component) and gaze traces (**Figure 3**). For this analysis, we utilized data from trials with complete visibility of the falling point (*i.e*., from trials without mask, mask type = 0). As expected, increasing the variability of the falling target trajectory resulted in an increased curvature in manual responses (linear regression model fit, m = 0.02, *P* < 0.001; *n* = 41; **Figure 3a**), but notably, ocular responses did not show such a rise in curvature (m = 0.14; *P* = 0.19; *n* = 20; **Figure 3b**).

**FIGURE 3.**
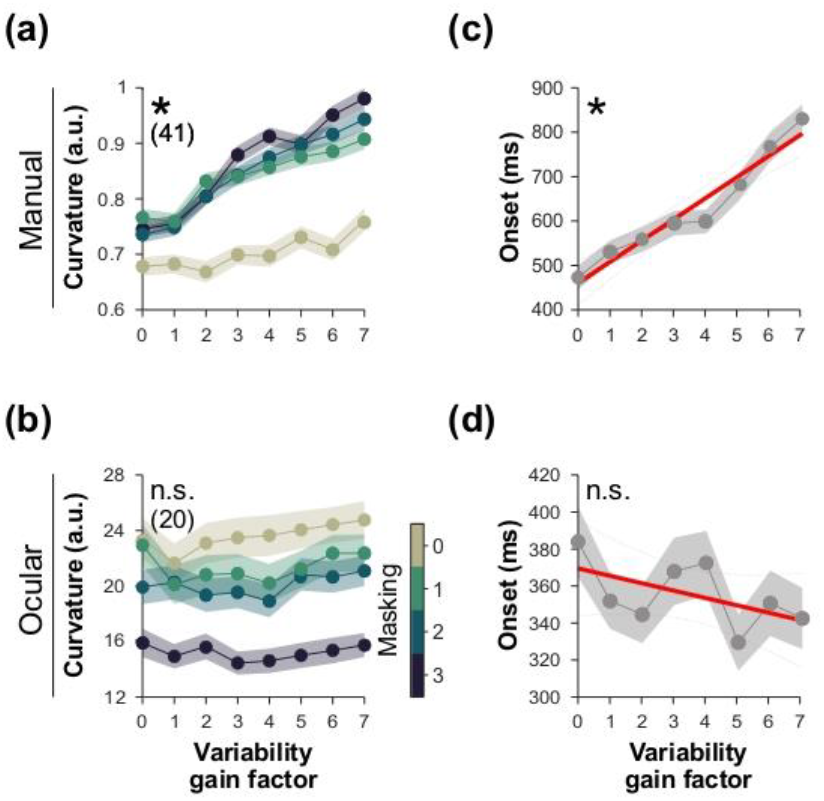
Curvature and onset of catching responses. (a) Curvature analysis of catching trajectories and (b) gaze data, presented with respect to target variability (x-axis) and masking conditions (colorbar). Predictive control is characterized by lower curvature and more linear chasing trajectories, while prospective control exhibits higher curvature with continuous adjustments. (c) Onset timing analysis (in ms) for manual responses and (d) gaze responses, with thresholds set at 30°/s for manual and 50°/s for ocular movements (see Methods). Participant numbers are indicated in parentheses. Asterisks depict significant relationships.

Another way to distinguish predictive from prospective control is to quantify the timing of the chaser’s movements (Brenner & Smeets, 2011; Fooken et al., 2016). In predictive control, the pursuer commonly initiates movement earlier than in prospective control. We established thresholds of 30°/s and 50°/s to determine the onset of manual and saccadic eye movements, respectively (Klein & Ettinger, 2019). The selection of a smaller threshold for horizontal manual movements was determined empirically by calculating the root mean square (RMS) of traces before and after movement initiation. Simultaneously, the chosen threshold for eye tracking data, validated in previous studies for distinguishing rapid from slower eye movements, ensured that only rapid and ‘ballistic’ ocular movements were included in the onset analysis (Klein & Ettinger, 2019). As mentioned, in predictive control, the chaser will tend to initiate their eye movements earlier than in prospective control. However, if the target’s trajectory changes more frequently, the chaser should need more time to initiate and adjust their course. In agreement, our analysis revealed that the falling object’s trajectory variability contributed to a notable increase in the onset of manual responses (m = 48.98; *P* < 0.0001; **Figure 3c**), but again, it did not change the onset of ocular responses in our task (m = -4.02; *P* = 0.16; **Figure 3d**). The negative slope from this regression may suggest that the ocular system tends to adjust to heightened variability by initiating responses slightly more rapidly. However, the main result is that target variability increased the onset latency of manual responses but did not show a similar effect on ocular responses.

### 3.4 Divergent dynamics between gaze and manual strategies

Visuomotor tasks require the coordination of visual and motor systems for intercepting or tracking moving targets. In simpler tasks with predictable target trajectories, gaze position is a reliable predictor of motor output, as the ocular system can accurately track the target using smooth pursuit eye movements (Fooken *et al*., 2016). Indeed, gaze position has been recognized as a robust predictor of manual response in interception tasks, where the eyes follow the object’s trajectory, improving interception accuracy (Land & McLeod, 2000). This phenomenon, known as ‘gaze-hand coupling’, indicates that people’s gaze position can predict the location of their hand at the interception point with relatively high accuracy (Goodale & Milner, 1992). However, the correspondence between gaze and manual response is not always perfect, with a consistent delay of approximately 50-100 milliseconds between gaze and hand movements during interception (Ricker *et al*., 1999; Bootsma *et al*., 2010). Moreover, the relationship between gaze position and manual response can vary across interception tasks (Bootsma *et al*., 2010).

Various factors could contribute to the coupling strength between gaze position and manual response, including the speed and direction of the moving object, the object’s distance, and the individual’s skill level. The ‘gaze-hand coupling’ associated with the visuomotor task we employed, characterized by high target variability, remains unknown. We analyzed the forward traces obtained from trials without a mask (*i.e*., occlusion level = 0). For each variability condition (target variability), we computed the area under the curve (AUC) for the respective forward traces (25 variants). Subsequently, we computed the sum of all these areas (ΣAUC) and assigned it to each participant. This cumulative value served as an integrated measure, characterizing the participant’s prototypical response (*i.e*., a single number leaning toward predictive or prospective). The ΣAUC, obtained from manual responses (*i.e*., output performance), was sorted across participants, ranging from the lowest to the highest values (left panel in **Figure 4a**). Maintaining consistent color tags for participants, **Figure 4b** illustrates that the ΣAUC, derived from gaze data, did not exhibit an organized relationship against the manual response (m = 3.8 × 10^4^; *P* = 0.97; *n* = 20). This finding gains further support when comparing the ΣAUC obtained from manual responses with those obtained from the gaze data of the same individuals (m = 0.04; *P* = 0.77; **Figure 4c**). Yet, this method may seem relatively complex, as it involves evaluating both predictive and prospective strategies based on ΣAUC of the forward traces (as illustrated in **Figure 2**).

**FIGURE 4.**
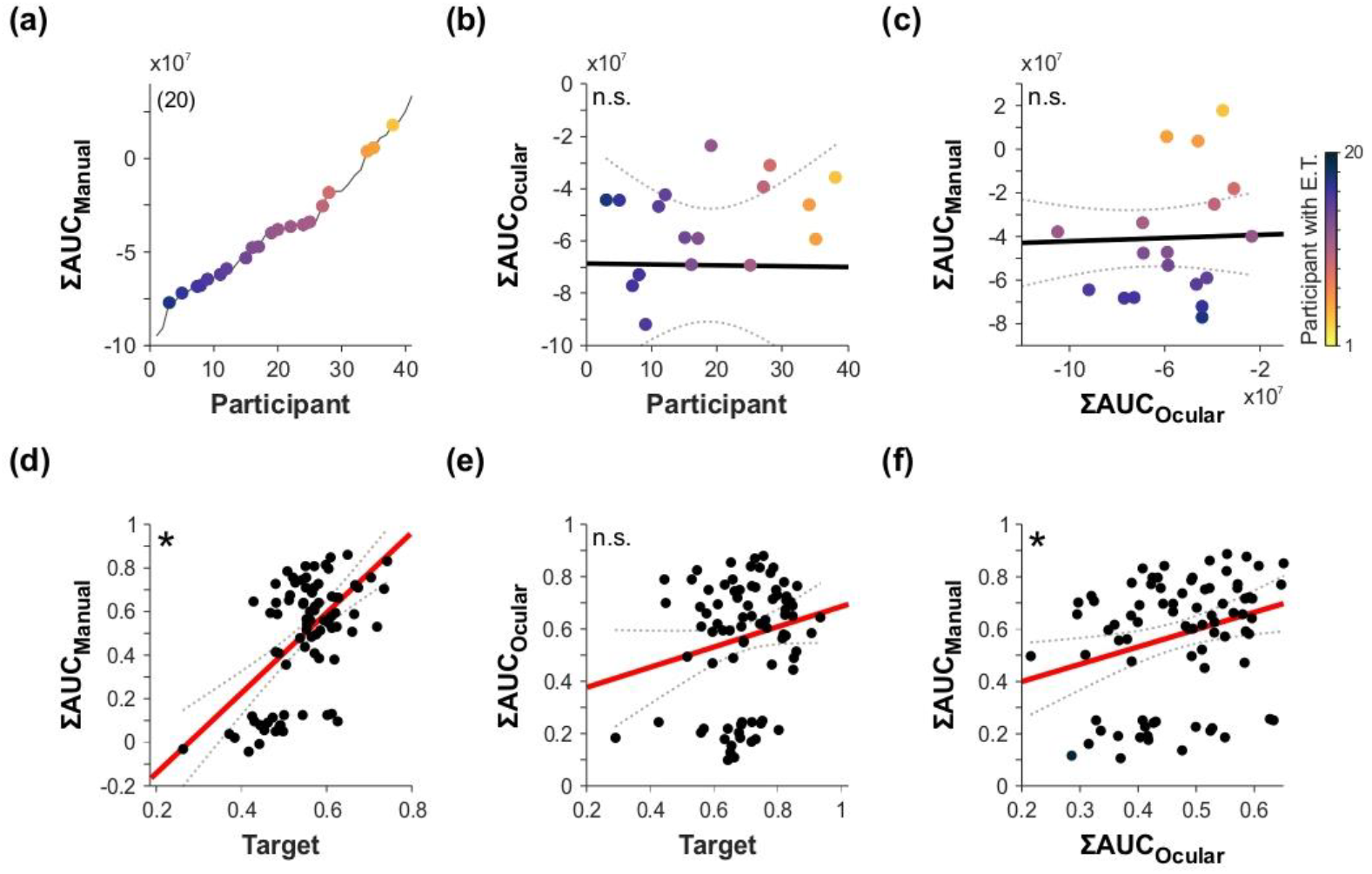
Gaze-hand coupling analysis for the falling dot task. The ΣAUC of forward traces was computed, representing each participant’s prototypical response. (a) ΣAUC relationships from manual responses sorted from lowest to highest, while (b) those from gaze data showed no apparent structure. (c) Limited correspondence between ΣAUC manual and gaze responses, maintaining color-coding for the dots across the first three panels. Cross-correlations among signals, including the moving target’s position, manual response, and gaze response. (d) Robust dependencies between manual response and target position, in contrast with (e) gaze response and target position, and a (f) weaker relationship between manual response and gaze response. Number of participants in parentheses. Asterisks depict significant relationships.

To approach this comparison differently, we computed cross-correlations across averaged recorded signals during each trial, such as the position of the moving target, the manual response, and the gaze response (**Figure 4d**). Subsequently, linear regressions and correlations were computed. We observed robust dependencies between manual response and target position (m = 1.62; *P* < 0.001; **Figure 4d**), and a comparatively weaker relationship between manual response and gaze response (m = 0.65; *P* < 0.02; **Figure 4f**). Moreover, gaze response did not predict target position (m = 0.39; *P* = 0.08; **Figure 4e**). Our results suggest that gaze position is a poor predictor of motor output in our task. This implies that the ocular system, probably engaged in active exploration, did not follow a smoothly predictable trajectory. Most likely, in our task, the ocular system explored the environment and gathered information beyond tracking the target. **Supplementary Figure 3** presents the analysis of gaze-hand coupling under diverse masking conditions.

### 3.5 Individual Variability in the Utilization of Predictive and Prospective Strategies

Previous studies showed the impact of traits and individuals’ choice biases on the strategies for solving some of our tasks (Trevino *et al*., 2020; Trevino *et al*., 2021). Examining individual differences is often addressed through test-retest experiments, which involve administering an identical test to the same cohort of participants at two distinct time intervals. The outcomes are then compared to quantify the stability of the scores. We reasoned that there could be individual differences in the reliance on predictive versus prospective control, depending on factors such as skill level, task demands, or personal preferences. To investigate the stability of strategies within a single session, we adopted a test-retest approach by segmenting the experimental session into three blocks. In the initial block (Test: baseline for comparison), participants undertook the main experiment, solving the task through permutated variability conditions and masks for the first 200 trials. The subsequent 200 trials (Intermediate phase) introduced a novel element: the color of the falling dot indicated the side (green or red) and initial horizontal speed of the ongoing trial. This visual information was intended to offer participants a clear cue regarding the falling trajectory, potentially shaping their utilization of predictive and prospective strategies. Finally, the last 200 trials (Retest) replicated the conditions of the initial phase. Data collected during task performance and testing were analyzed to identify individual differences using correlations. Comparative analyses between the second and first epochs and between the third and first epochs revealed high correlations for both comparisons (test-retest procedure within a single session; each gray dot represents the ΣAUC across variability factors per participant; Pearson correlation coefficient ≥ 0.80, *P* < 0.005; *n* = 10; **Figure 5a**), indicating good reliability.

**FIGURE 5.**
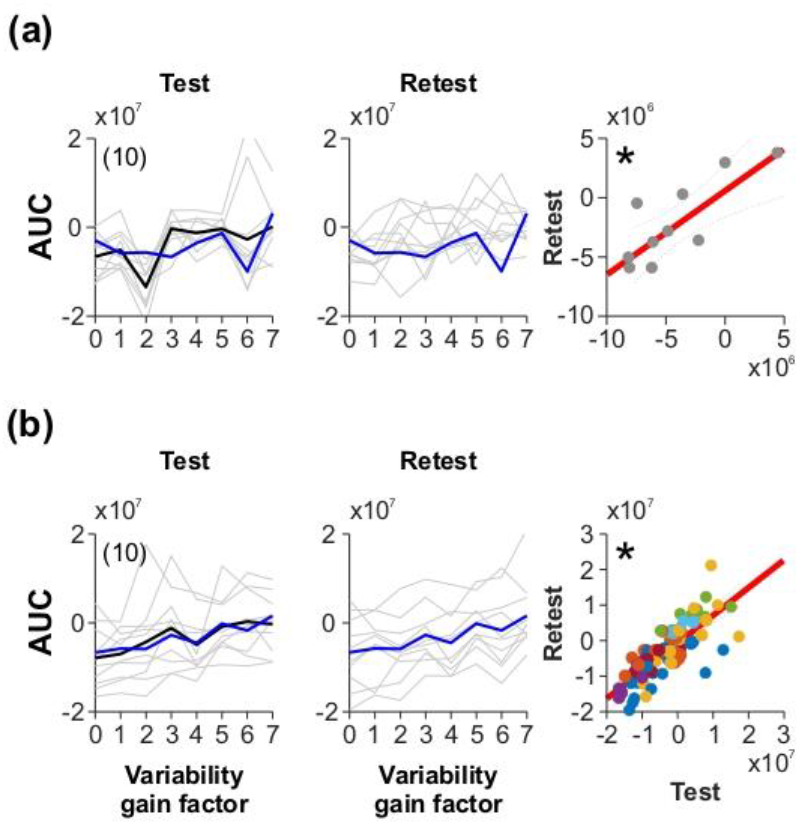
Stability of predictive and prospective strategies. (a) Test-retest procedure within a single session. Participants engaged in the test phase where they performed the main experiment (left panel). An intermediate phase introduced a color rule for trajectory anticipation (not illustrated, see Methods), and the retest phase replicated the initial conditions (middle panel). High correlation between the third and first epochs indicate test-retest reliability (right panel). (b) In a separate experiment, a new group of participants underwent a two-week interval test-retest, utilizing their ΣAUC of manual responses (color-coded for the different variability levels), revealing satisfactory reliability and consistency of predictive/prospective control strategies. Number of participants in parentheses. Asterisks depict significant relationships.

We subsequently investigated whether a two-week interval between tests in a test-retest comparison would yield similarly high correlations. This examination involved a new group of participants, with the results indicating satisfactory reliability (AUC values for variability conditions are represented using consistent, randomly assigned colors for each participant; Pearson correlation coefficient = 0.78; *P* < 0.0001; *n* = 10; **Figure 5b**).

## 4 DISCUSSION

Interception is a fundamental visuomotor skill in everyday activities like driving and sports. Real-world situations present dynamic and complex environments where multiple interception strategies may come into play. Here, we investigated how predictive and prospective interception strategies dynamically interact. We designed a visuomotor task, systematically manipulating target trajectory and visibility through external forces and graded masking techniques. We characterized their interception strategies by analyzing the participants’ manual and gaze responses under controlled conditions. Our results revealed that participants employed a combination of predictive and prospective strategies when intercepting a falling target on a screen, with the choice between these strategies contingent on the target’s predictability and visibility. Masking yielded an interesting outcome, diminishing the predictive component more prominently in manual traces than in gaze data, thereby pushing the interception strategies toward greater dependence on a prospective approach. In manual responses, a shift from predictive to prospective strategies occurred in situations with increased unpredictability and reduced visibility of the target’s motion, contrasting with the preference for predictive strategies in scenarios marked by higher predictability. This shift in strategy was paralleled by the increased curvature of hand trajectories, indicating continuous adjustments made to intercept the target under our varying conditions.

Predictive strategies rely on anticipating a moving target’s future position through mechanisms like forward modeling and predictive processing (Clark, 2023). Although relatively easy to implement, purely predictive interception may encounter difficulties in dynamic environments. Alternatively, prospective strategies involve real-time monitoring and rapid adjustments, demanding effective hand-eye coordination and sensorimotor integration. Thus, despite being more difficult to implement, prospective control offers greater flexibility, especially in complex interception scenarios. In principle, prospective strategies may require additional neural resources, engaging in complex cognitive processes such as prediction, decision-making, and adaptive control. The relative difficulty between the two strategies is context-dependent and varies based on the specific requirements of the interception task. Both strategies have unique challenges and advantages, allowing for complementary use depending on the circumstances.

The study of predictive and prospective control in dynamic tasks suggests that there could be a sequential utilization of these strategies at different stages of the interception process. More specifically, individuals may initiate prospective interception to approach the target but then transition into predictive control as the optimal moment for collision approaches (Treviño *et al*., 2023). Some authors argue that information-based prospective control is more parsimonious and plausible than preprogrammed timing (Bootsma *et al*., 2010). Some others propose that interception can be solved with a preprogrammed timing approach, which implies that the movement is initiated and executed without major visual corrections (van Soest *et al*., 2010). Our findings imply that a viable solution may lie somewhere in the middle. Indeed, some practical applications suggest an interchangeable use of these strategies, enhancing flexibility in interceptive actions (Hodges *et al*., 2004).

We recently explored how people chase moving targets with different speeds and directional uncertainty levels on a screen. We introduced a geometric model to quantify and differentiate pursuit and interception strategies based on the optimal angles for each behavior. The analysis revealed that the predictive interception strategy increased as the collision moment approached and was influenced by the target’s speed, direction, and visibility (Treviño *et al*., 2023). We also found that gaze behavior was adjusted by the target’s position and velocity, indicating a prospective control of interception. Here, we found that reduced target variability favored predictive strategies, while increased variability prompted prospective strategies. Our results reveal the simultaneous utilization of predictive and prospective interception strategies in our experimental setup, with notable consistency observed across manual responses. Hence, our findings demonstrate that predictive and prospective interception strategies can coexist within the same task.

One way to study predictive and prospective control is through computer simulations (van Soest *et al*., 2010). Another way to address interceptive control is by exploring the chaser’s ocular movements. In predictive control, the chaser’s eye movements tend to be more anticipatory, as they constantly track the target’s predicted location. In prospective control, the chaser’s eye movements are more reactive, as they respond and track the target’s current location. Contrary to the mixed strategies we found in the manual responses, our analysis of gaze data indicates a prevalence of predictive strategies in ocular responses. It’s crucial to highlight that the positive differences observed in the manual response traces, indicative of a prospective component, do not necessarily imply the absence of a predictive element, as evidenced in the gaze data. Interestingly, a previous work investigated how people intercept moving targets on a screen with different levels of predictability. The authors measured the gaze, head, and hand movements in three experiments with different target trajectories and hitting zones. They showed that knowing the interception location in advance leads to faster, more direct, and more accurate movements than not knowing it. Thus, these results indicate that the predictability of the interception location is more important than the predictability of the target motion for interceptive actions (de la Malla *et al*., 2019).

We also examined the curvature and timing of the manual and gaze responses to distinguish between predictive and prospective control. In prospective control, the trajectory typically exhibits higher curvature as the chaser continually adjusts its course to intercept the erratically moving target. Contrastingly, in predictive control, the trajectory tends to be more linear, reflecting the chaser’s straightforward movement toward the target’s predicted location (*i.e*., less curvature). Analyzing the curvature of manual responses, we noted an increase with higher target variability, indicating a shift from predictive to prospective control. This adaptation suggests that constant adjustments are necessary in unpredictable conditions, leading to more curved manual trajectories. Interestingly, the eye trajectories did not exhibit a similar curvature increase with the stimulus trajectory’s unpredictability. This agrees with the fact that gaze position was a poor predictor of hand position in the task, suggesting that the eye movements were not simply following the target’s trajectory but instead exploring the environment and adapting to the variability of the task.

Another distinguishing factor lies in the timing of movements. In predictive control, the chaser initiates movement earlier than in prospective control. This early initiation allows the chaser to accommodate potential changes in the target’s trajectory, emphasizing the proactive nature of predictive control compared to the more reactive characteristics of prospective responses. Examining movement initiation, we found that predictive involved earlier onsets than prospective control. This was reflected in manual responses, with their onset delayed with higher target variability. In contrast, the rapid onset of gaze responses in unpredictable conditions suggests an adaptive mechanism to gather information efficiently. These findings highlight the dynamic interaction between predictive and prospective control and how the visual and motor systems adapt to meet task demands.

Research on the interplay between gaze position and motor output in complex visuomotor tasks has yielded inconsistent findings (Land & McLeod, 2000; Henderson, 2017). Some studies assert that gaze position is a crucial predictor of motor output, while others challenge this association. The outcome variability highlights the intricate and non-linear relationship between gaze position and motor output, dependent on task specificity, individual proficiency, and environmental demands. Our examination of hand-eye movements found that the coordination between eye movements and manual performance had a low coupling. This might be attributed to the eyes being engaged in additional ‘visual scanning’ tasks, diminishing their efficiency as predictors, particularly under more variable target conditions. Thus, the premise is that the gaze was engaged in various activities beyond task resolution, mainly triggered by external variability. Consequently, it became a poor predictor of the manual output.

The stability of participants’ interception strategies was evident within a single session and over a two-week interval, indicating robust reliability in their preference for predictive or prospective strategies. Some participants consistently leaned towards predictive, while others favored prospective strategies, and these preferences remained consistent over time. Therefore, it is essential to acknowledge that these are individual differences. The coexistence of predictive and prospective strategies opens the door for developing tailored training programs that leverage these individual differences, enhancing performance in visuomotor tasks across diverse populations.

Our study has important limitations that should be considered. Firstly, this initial investigation did not address the neural mechanisms underlying the shift between predictive and prospective strategies, creating a gap in our understanding of the associated neurobiological processes. Additionally, the role of individual factors, including experience, training, and cognitive abilities, remains insufficiently examined. For example, our study lacked assessments of cognitive load, attention, and motivation, pivotal elements that could significantly influence participants’ performance and strategy selection (Parr & Friston, 2017; Ouwehand *et al*., 2021). Therefore, although recognized as important variables determining interception performance, these factors’ cumulative impact is yet to be fully elucidated. Finally, the computer-based nature of our task raises concerns regarding ecological validity, as it may not entirely replicate real-world interception scenarios like those encountered in sports or driving. Future research should aim to address these limitations for more practical applications.

## 5 CONCLUSION

Our results highlight the coexistence of predictive and prospective strategies in a visuomotor task, where participants catch a falling dot on a screen. The study demonstrates that the balance between these strategies depends on factors like target predictability and visibility, with reduced variability favoring predictive strategies and increased variability promoting prospective ones. The analysis of manual and gaze responses reveals distinct adaptations, indicating that target characteristics significantly impact manual responses, whereas gaze responses display a more predictive and exploratory nature. The task, centered on catching the target through manual control of an optical mouse, imposing no requirements on how participants should utilize their gaze.

Experimentally controlled factors substantially influenced the shift from predictive to prospective strategies. However, the role of individual differences and internal factors should not be overlooked. This means that cognitive processes may be crucial in allocating internal resources to implement predictive or prospective strategies (Treviño *et al*., 2023). Can individuals be trained to control the relative utilization of these two strategies consciously? What would be the potential advantages or disadvantages associated with such volitional control? Understanding when and how individuals switch between predictive and prospective strategies holds the potential for developing targeted training methods. Applications in sports, gaming, and tasks demanding rapid decision-making based on predictions could benefit from enhanced strategy-switching abilities. Beyond practical domains, this understanding may have clinical implications, particularly in conditions affecting the utilization of these strategies (Clark, 2023). Insights into human strategy use can inform the design of artificial systems, contributing to developing more effective and responsive technologies.

## AUTHOR CONTRIBUTIONS

*Conceptualization:* MT; *Methodology:* MT, IM; *Validation:* IM; *Formal Analysis:* MT; *Investigation:* IM; *Data curation and formal analysis:* MT; *Funding acquisition:* MT, IM; *Visualization:* MT, IM; *Writing-original draft:* MT; *Writing-review and editing:* MT.

## DATA AVAILABILITY STATEMENT

Upon a reasonable request, interested parties can obtain the datasets generated in this study by contacting the corresponding author.

**SUPPLEMENTARY FIGURE 1.**
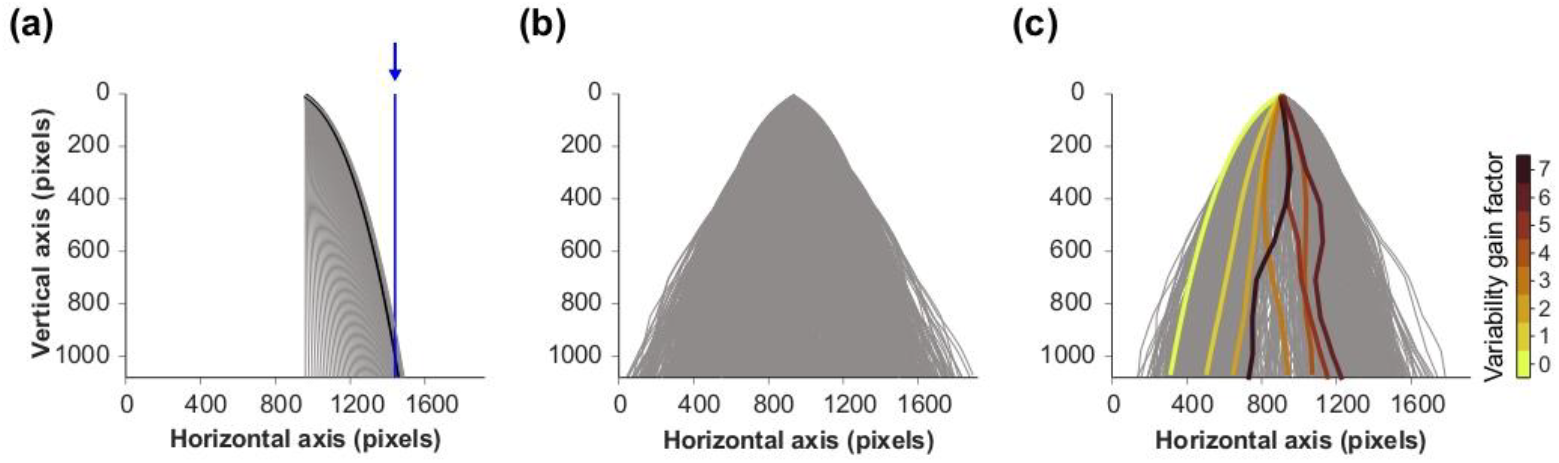
Experimental Task Design. The panels depict the sequential steps involved in designing and implementing the experimental task. (a) The initial phase involved simulating the trajectory of the falling point and adjusting parameters such as initial speed, friction acceleration, and falling acceleration to achieve a consistent descent to the midpoint of the monitor screen’s left or right half within a 2.3-second trial duration. (b) To introduce variability, six additional forces were strategically applied along equidistant imaginary lines and intermediate heights, perturbing the horizontal trajectory of the falling particle, and influencing its speed and landing position. The direction and intensity of these forces were deliberately adjustable to create varying degrees of impact on the particle’s path. Computer simulations were conducted to identify force magnitudes and directions, and a combination of random forces was selected to ensure final target positions within the screen boundaries. (c) Finally, a virtual gain factor, adjustable from 0 to 1, was introduced, allowing systematic and precise modifications to the impact of these external forces.

**SUPPLEMENTARY FIGURE 2.**
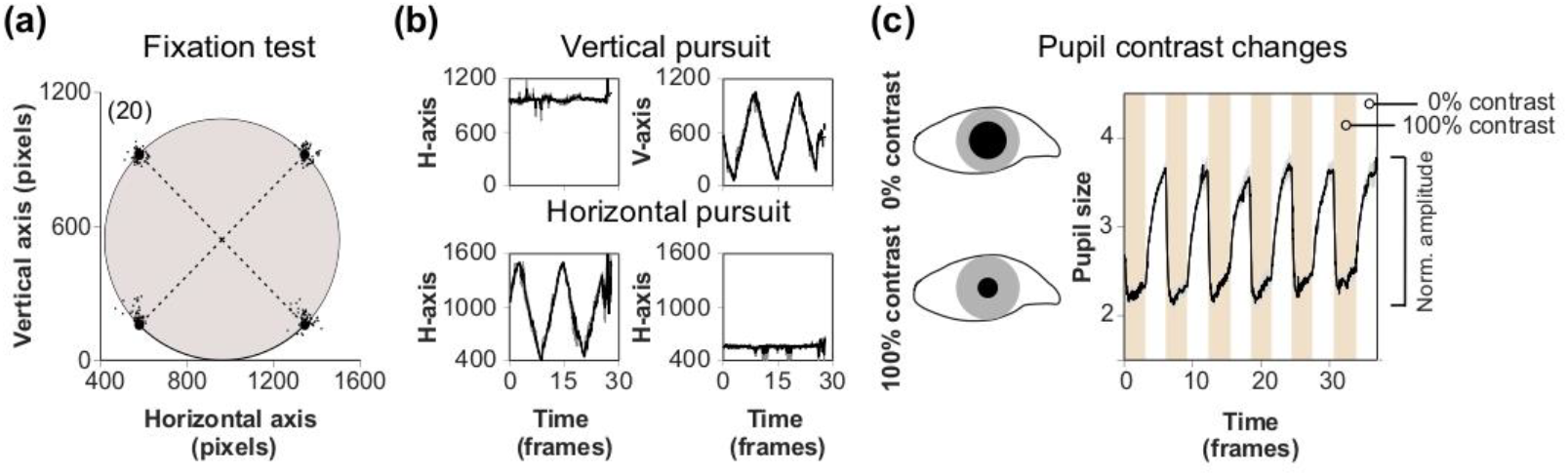
Eye-tracker calibration. (a) The calibration involved a four-point routine with fixation points strategically positioned around the periphery of a circle centered on the screen, matching its diameter to the height of the monitor. (b) Following the routine, participants engaged in a smooth pursuit task, tracking a white dot’s horizontal or vertical movement along the same circle at a speed of 10 °/s. (c) The final calibration step assessed changes in pupil size as participants transitioned between a very bright and a completely black screen. The right panel displays the average pupillary diameters in response to rapid changes in screen contrast from 0 to 100%. Individual participant responses were recorded for calibration of subsequent pupillary measurements. Number of participants in parentheses.

**SUPPLEMENTARY FIGURE 3.**
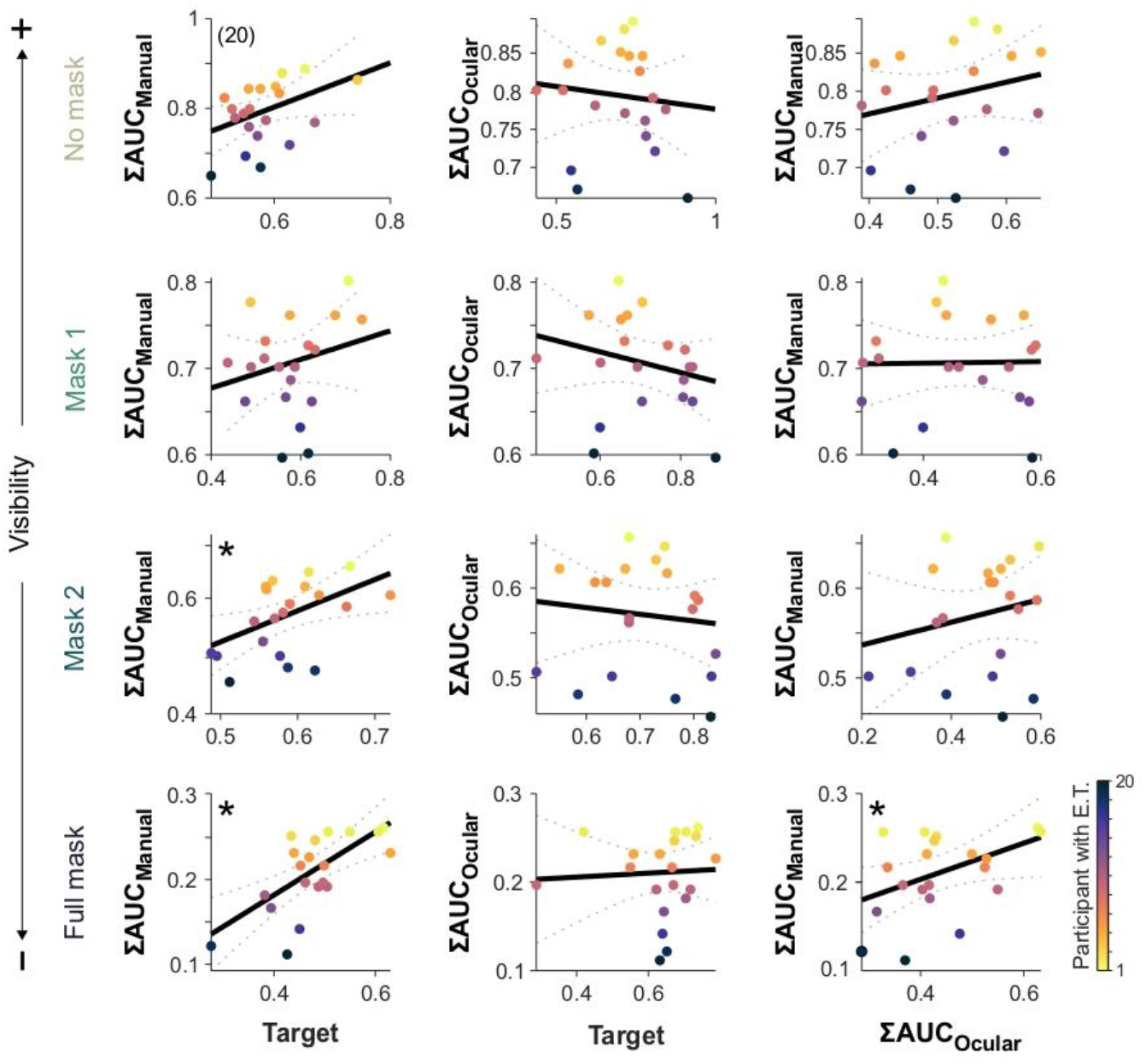
Gaze-Hand coupling analysis across masking conditions. This figure expands upon the gaze-hand coupling analysis presented in Figure 4, specifically investigating the impact of various masking conditions on visuomotor task performance. Summed area under the curve (ΣAUC) on forward traces was employed, with consistent color-coding for participants, sorted from lowest to highest ΣAUC. Columns illustrate comparisons between manual responses and the moving target’s position (left), gaze responses and the moving target’s position (middle), and manual responses versus gaze responses (right). Rows depict different masking conditions, ranging from mask type 0 (higher visibility, upper row) to mask type 3 (lower visibility, lower row). Number of participants in parentheses. Asterisks depict significant relationships.

## Acknowledgements

Thanks to I. Sevilla for helping to conduct some experiments. Thanks to Coordinación de Investigación, Secretaría Administrativa, Secretaría Académica, and the leadership and professionalism from both of our university centers (CUCBA, and CUCIÉNEGA), and Instituto de Neurociencias for their constant support. We thank Dr. J. Yau (Baylor College of Medicine) for critical reading and insightful suggestions. We appreciate the time the reviewers dedicated to helping us improve our manuscript.

